# Deletion of Cd248 in *Postn* positive myofibroblast Fails to Attenuate Cardiac Remodeling and Fibrosis in a Murine Model of Pressure-Overload Cardiomyopathy

**DOI:** 10.1101/2025.09.26.678917

**Authors:** Donghua Li, Dongmei Zhong, Tian Tan, Zhilei Huang, Mingyue Wu, Yongshan Liu, Yalin Zhang, Chen Liu, Fu-Li Xiang, Jie Xu

## Abstract

**Background:** Cardiac fibrosis, driven by activated Periostin (Postn)-expressing myofibroblasts, is a key pathophysiological feature of heart failure induced by pressure overload. The clinic trial tested tumor vascular regulator CD248 has been validated as a potential anti-fibrotic target in ischemic heart injury. We therefore tested the hypothesis that targeting Cd248 specifically in *Postn*^+^ myofibroblasts ameliorates pressure-overload cardiomyopathy.

**Methods:** Single-cell RNA-sequencing data from murine hearts subjected to transverse aortic constriction (TAC) were analyzed to quantify the *Cd248*^+^*/Postn*^+^ fibroblast population. Subsequently, *Postn*^+^ myofibroblast-specific inducible *Cd248* knockout mice were subjected to TAC. Gene deletion was induced with tamoxifen from the day of surgery. Cardiac remodeling and function were analyzed 8 weeks after TAC.

**Results:** Analysis of scRNAseq data confirmed that about 1/3 of the fibroblasts in the 14-day TAC heart are *Cd248*^+^*Postn*^+^ fibroblasts. However, comprehensive echocardiographic analysis revealed no significant differences in key parameters of cardiac function or dimensions, including left ventricular ejection fraction, ventricular volumes, wall thickness and global muscle strain between *PostnMCM*+*/-;Cd248fl/fl* and *PostnMCM*+*/-* controls. Histological analysis at 8 weeks also showed no significant alteration in the extent of interstitial or perivascular fibrosis, myocardial vessel density, or cardiomyocyte hypertrophy.

**Conclusion:** The specific deletion of Cd248 in *Postn*-expressing myofibroblasts does not mitigate pathological remodeling or fibrosis in response to chronic pressure overload. These findings suggest the role of Cd248 in this cell lineage is not a primary driver of pressure-overload pathology. Therefore, a better understanding of the context-dependent functions of CD248 is critical to successfully guide the clinical translation of CD248-based anti-fibrotic therapies.

## Introduction

Heart failure, a leading cause of mortality worldwide, often results from maladaptive cardiac remodeling in response to chronic injury stimuli such as pressure overload from hypertension or aortic stenosis^[1]^. A central pathological hallmark of this process is cardiac fibrosis, characterized by the excessive deposition of extracellular matrix (ECM) proteins^[2]^. While initially a reparative response, sustained fibrosis leads to increased myocardial stiffness, diastolic dysfunction, and arrhythmogenic substrates, ultimately driving the progression to heart failure. Therefore, elucidating the cellular and molecular drivers of fibrosis is urgently needed for developing effective new therapies.

The resident cardiac fibroblast (CF) has emerged as the principal architect of both adaptive and maladaptive cardiac remodeling^[2]^. Upon injury, quiescent CFs activate and differentiate into myofibroblasts, which are characterized by the expression of proteins such as Periostin (*Postn*) and smooth muscle actin (*SMA*)^[3]^. As SMA also strongly expressed in smooth muscle cells, the *Postn*-Cre and *Postn*-MerCreMer driver became invaluable and widely used tools for specifically targeting activated CF population *in vivo*^[2, 4, 5]^. *Postn*^+^ fibroblast-specific deletion of the transcription factor KLF5 was shown to attenuate not only pressure overload-induced fibrosis but also cardiomyocyte hypertrophy, revealing a critical paracrine cross-talk (via IGF-1) between fibroblasts and muscle cells^[6]^. Similarly, our own work ()and that of others have demonstrated that targeting the canonical WNT/β-catenin or TGF-β-Smad3 signaling pathway specifically within the *Postn*^+^ fibroblast lineage is sufficient to robustly block the fibrotic response to pressure overload^[3, 7]^. Moreover, the specific ablation of the collagen chaperone Hsp47 in *Postn*^+^ fibroblasts abrogated pressure overload-induced fibrosis and blunted the hypertrophic response, confirming *Postn*^+^ CFs as the primary source of pathological collagen^[8]^. These studies solidify the *Postn*^+^ CFs as a central and validated target for anti-fibrotic therapies.

Cd248 is a type-I transmembrane glycoprotein with C-type lectin and EGF-like domains that is expressed at low levels in most adult tissues but is induced on mesenchymal stromal cells—notably pericytes, vascular smooth muscle cells (VSMC) and fibroblasts—during development, inflammation and remodeling^[9, 10]^. Genetic and biochemical studies show that Cd248 participates in vascular patterning and plays a modulatory role in microvascular remodeling^[11]^. Cd248 promotes VSMC remodeling and plaque development in atherosclerosis^[12]^. Recently, two parallel studies using single-cell transcriptomics discovered that Cd248 (Endosialin/TEM1) is a specific surface marker for a late-activated, pathological fibroblast subpopulation that drives chronic fibrosis after myocardial infarction (MI) or ischemia/reperfusion (I/R)^[13, 14]^. Critically, both studies demonstrated that therapeutic interventions targeting these Cd248^+^ cells—using either CAR-T cell therapy or neutralizing antibodies—successfully ameliorated cardiac fibrosis and improved heart function. One of these studies further showed that deleting *Cd248* specifically in the *Postn*+ lineage was sufficient to confer this protection in an I/R model^[14]^.

However, whether Cd248 is required for fibrosis in pressure-overload cardiomyopathy is unknown. Here, motivated by an siRNA screen in primary cardiac fibroblasts that highlighted Cd248 as a regulator of TGF-β–driven activation, we tested the hypothesis that selective deletion of Cd248 in Postn^+^ myofibroblasts improves pathological remodeling in pressure-overload cardiomyopathy induced by transverse aortic constriction (TAC). Contrary to expectation, myofibroblast-restricted Cd248 deletion did not reduce TAC-induced fibrosis or pathological remodeling, despite clear cell-autonomous effects of CD248 loss on fibroblast activation, migration, and proliferation in vitro. These findings suggest that CD248’s role in cardiac fibrosis is context-dependent, differing across injury types and disease stages, and they emphasize the need to match fibroblast-targeted therapies to the appropriate pathological niche.

## Methods

### Animal and Husbandry

All animal experiments were approved by the Sun Yat-sen University Animal Care and Use Committee (SYSU-IACUC-2024-002343). Myofibroblast-specific inducible *Cd248* knockout mice (*PostnMCM*+*/-;Cd248fl/fl*) and their corresponding controls (*PostnMCM*+*/-*, The Jackson Laboratory, 029645) were generated on a C57BL/6J background. To validate Cre recombinase activity, *PostnMCM*+*/-* mice were crossed with a TdTomato reporter line containing a Loxp-Stop-LoxP cassette. All mice were housed in a specific pathogen-free (SPF) facility on a 12-hour light/dark cycle with ad libitum access to food and water. All surgical and experimental procedures were performed on male mice aged 10-14 weeks.

### Experimental Design and SAGER Guidelines

In accordance with SAGER (Sex and Gender Equity in Research) guidelines, we report that this study was conducted exclusively in male mice aged 10-14 weeks. This single-sex design was chosen to minimize biological variability associated with the estrous cycle in females and to avoid the known confounding cardioprotective effects of female sex hormones in cardiac injury models, thereby reducing the total number of animals required to achieve statistical power. We acknowledge that this approach limits the direct generalizability of our findings. The role of Cd248 in pressure-overload cardiomyopathy may differ in females, and future studies are warranted to investigate potential sex-specific mechanisms.

For comparisons between genotypes, littermate controls were used whenever possible. Within each genotype, mice were randomly allocated to either the transverse aortic constriction (TAC) or sham surgery group. Key outcome assessments, including the analysis of echocardiographic data and the quantification of histological fibrosis and cardiomyocyte size, were performed by investigators who were blinded to the genotype and surgical group of the animals.

### Statistical Analysis

All quantitative data are presented as mean ± SEM. Statistical analyses were performed using GraphPad Prism software (v9.0, GraphPad Software). For comparisons between two groups with normally distributed data, an unpaired, two-tailed Student’s t-test was used. For comparisons between more than two groups with normally distributed data, a one-way or two-way Analysis of Variance (ANOVA) was performed, followed by Tukey’s multiple comparisons post-hoc test. For data that were not normally distributed, the non-parametric Mann-Whitney U test (for two groups) or the Kruskal-Wallis test with Dunn’s multiple comparisons test (for multiple groups) was used. Survival curves were generated using the Kaplan-Meier method and compared using the Log-rank (Mantel-Cox) test. A P-value < 0.05 was considered statistically significant. The specific tests used for each experiment are detailed in the figure legends.

Please see detailed methodology in **Supplemental method**.

## Results

### Single-cell analysis reveals Cd248 expression in activated cardiac fibroblasts

To confirm CD248 expression in fibroblast in pressure overload disease setting, we analyzed published mouse and human single-cell RNA-sequencing datasets. In a public dataset from mouse hearts subjected to TAC (GSE166403)^[15]^, the *Cd248*-positive cell population expanded (Figure 1A and B), with expression predominantly in fibroblasts, followed by mural cells and endothelial cells (Figure 1B). Within the fibroblast population 14 days after TAC, the number of *Postn*^+^ cells increased dramatically (Figure 1A and C), and the majority of *Cd248*^+^ cells co-localized with *Postn* resulting in about 33% of the total fibroblast being *Postn*^+^*Cd248*^+^ (Figure 1D). In a published human hypertrophic cardiomyopathy single nuclear RNAseq dataset^[16]^, CD248 was found to be significantly increased and majorly expressed by pericytes, fibroblasts and smooth muscle cells in hypertrophic human heart (Supplemental Figure 1A). This confirms our genetic strategy targets the intended pathological *Postn*^+^*Cd248*^+^ myofibroblast population.

**Figure 1.**
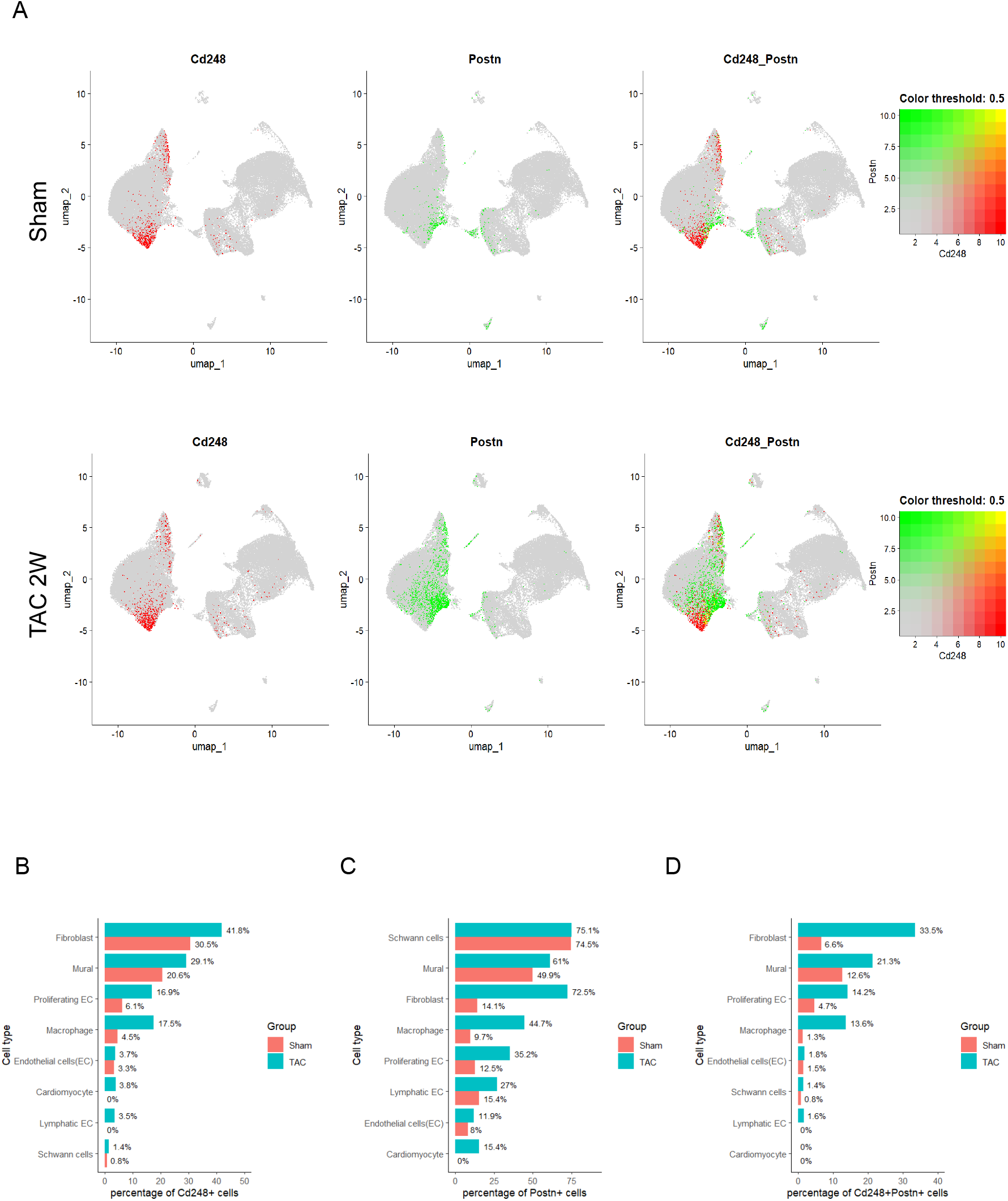
Single-cell RNA-seq reveals co-expression of Cd248 and Postn in cardiac fibroblasts and mural cells following pressure overload. (A, B) Uniform Manifold Approximation and Projection (UMAP) of cardiac interstitial cells from sham and TAC-operated mice, annotated by cell type (A) and experimental group (B). (C-H) Feature plots illustrating the expression of Cd248 (red), Postn (green), and their co-expression (yellow overlay) in hearts from sham (C-E) and TAC (F-H) groups. Insects show the bivariate color key; cells were considered positive if the normalized expression value exceeded 0.5. (I-K) Quantification of the percentage of cells positive for Cd248, Postn, or both within each major cell lineage. Data are presented as percentages for each group (salmon = Sham, teal = TAC).

We also validated *Cd248* gene expression in various organs and different cardiac injury models. We found that the gene expression of Cd248 is relatively high in heart, lung, kidney, skeletal muscle and brain (Supplemental Figure 1A). In TAC model, cardiac expression of Cd248 was significantly upregulated 1 week after TAC (Supplemental Figure 2A). The fibrosis marker (*Col1a1, Col3a1, Postn*, and *Acta2*) and mechano-stress associated membrane frangibility regulator *Ninj1* (Supplemental Figure 2B) also significantly increased. Moreover, consistent with published studies^[13, 14]^, upregulation of CD248 was also observed in the infarct border zone 3 days after MI (Supplemental Figure 2C).

### Cd248 knockdown attenuates myofibroblast activation, migration, and proliferation *in vitro*

To first establish the functional role of Cd248 in cardiac fibroblasts, we performed knockdown experiments in primary adult mouse cardiac fibroblasts (mCFs). Pathological stimulation with TGF-β1 robustly induced mCF activation into myofibroblasts, characterized by a ∼4-fold increase in *Acta2* (α-SMA) mRNA and the formation of prominent α-SMA stress fibers (Figure 2A and B). Knockdown of *Cd248* (about 80% efficiency, Supplemental Figure 3A) significantly inhibited this activation process (Figure 2A and B). Migration was measured by wound scratch assay. Knocking down of *Cd248* in mCF exhibited significantly slower wound healing compared to the control group (Figure 2C and D). Moreover, cell proliferation was detected by EdU staining. Both the total cell number and the number of proliferating cells marked by EdU were significantly lower in the TGF-β1 treated *Cd248* knockdown mCF (Figure 2E-G). *Cd248* knockdown mCF also showed significantly less *Col3a1* gene expression, but no decreases in *Col1a1* and *Postn* (Supplemental Figure 3B-D). These data suggested that *Cd248* might play an important role in mCF activation, migration and proliferation.

**Figure 2.**
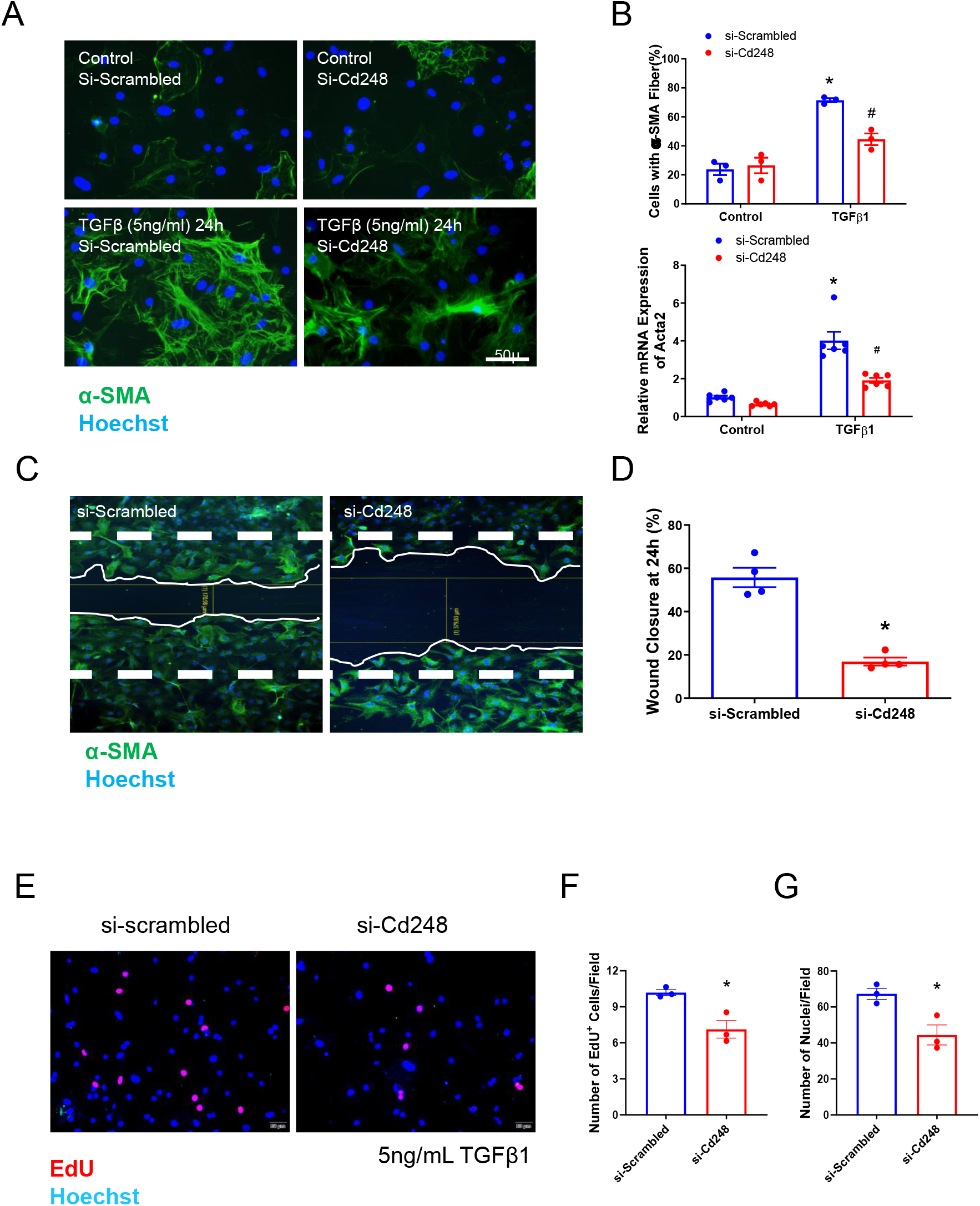
Knockdown of Cd248 blunts TGF-β1-induced fibroblast activation, migration, and proliferation in vitro. (A) Representative immunofluorescence of cultured cardiac fibroblasts transfected with control siRNA (si-Scrambled) or siRNA targeting Cd248 (si-Cd248) and stimulated with vehicle or TGF-β1 (5 ng/mL) for 24 hours. Myofibroblast activation is indicated by the formation of α-SMA stress fibers (green); nuclei are counterstained with Hoechst (blue). Scale bar, 50 µm. (B) Quantification of myofibroblast activation, showing the percentage of cells with organized α-SMA stress fibers and the relative mRNA expression of Acta2 (α-SMA). Two-way Analysis of Variance (ANOVA) was performed, followed by Tukey’s multiple comparisons post-hoc test. \**P* < 0.05 vs. the corresponding vehicle control; #*P* < 0.05 vs. si-Scrambled with TGF-β1 stimulation. (D) Scratch-wound assay demonstrating impaired fibroblast migration following Cd248 knockdown. Quantification shows the percentage of wound closure at 24 hours relative to the initial scratch. \**P* < 0.05 vs.si-Scrambled, determined by an unpaired Student’s *t-test*. (E-G) Analysis of fibroblast proliferation via EdU incorporation. Representative images (E) and quantification of the number of EdU^+^ nuclei (F) and total nuclei (G) per field. \**P* < 0.05 vs.si-Scrambled, determined by an unpaired Student’s *t-test*. n=3-6, data are presented as mean ± SEM; each dot represents an independent experiment.

### *Postn*^+^ myofibroblast-specific deletion of Cd248 has no impact on cardiac fibrosis or adverse remodeling in mouse TAC model

Given the pro-fibrotic function of Cd248 *in vitro*, we tested if deleting it in *Postn*^+^ myofibroblasts would protect against TAC-induced pathology. We validated the successful establishment of TAC model in house. In all TAC mice in this study, pulsed-wave Doppler echocardiography was used to confirm successful constriction, with peak blood flow velocity across the aortic arch increasing from a baseline of ∼1000 mm/s to at least 3500 mm/s post-TAC (Supplemental Figure 4A). Histological analysis via Picro-Sirius Red staining of WT hearts 6 weeks post-TAC confirmed the development of pathological fibrosis, showing a significant increase in both interstitial and perivascular fibrosis (Supplemental Figure 4B).

To specifically delete Cd248 in *Postn*^+^ myofibroblasts, we used the *PostnMCM*+*/-;Cd248fl/fl* (Postn-Cd248) and *PostnMCM*+*/-* (control) mouse lines, where tamoxifen induces Cre expression in emerging *Postn*^+^ cells. We validated this system using *PostnMCM*+*/-;TdTomato* reporter mice, which showed a progressive increase in TdTomato-positive myofibroblasts over time following TAC (Supplemental Figure 4C). The Cre recombination efficiency is comparable to the previous study using the same *Postn*-MCM system generated by Dr Mokentin^[3, 7]^.

Eight weeks after TAC, hearts from Postn-Cd248 knockout and control mice were harvested and submitted to comprehensive cardiac function and histology analysis. Peak blood flow velocity across the aortic arch in both groups was confirmed to be similar (Supplemental Figure 5A). Both groups have survival rates above 80% during the 8 weeks after TAC (Supplemental Figure 5B). HE staining of the 4-chamber view showed no changes in global morphology (Figure 3A). Quantification of Picro-Sirius Red-stained heart sections revealed that the fibrotic area, measured as both interstitial and perivascular collagen deposition, was extensive and statistically indistinguishable between the Postn-Cd248 knockout and control mice (Figure 3B and C). Furthermore, vessel density and myocardial hypertrophy were also analyzed. There were no significant differences in myocardial capillary density, cardiomyocyte cross-sectional area, or ventricular weight to body weight ratio between the two groups (Figure 4A-D). Cd248 has previously been shown to regulate arteriole remodeling in cancer ^[11, 17]^. Thus, we quantified the density of arteriole with three different size categories: <50μm, 50-100μm and >100μm. No difference in arteriole densities was detected between the Postn-Cd248 knockout and control mouse hearts (Figure 4E). Taken together, these histological data unequivocally demonstrate that deleting Cd248 specifically within the *Postn*^+^ myofibroblast lineage fails to mitigate the myocardial pathological remodeling of chronic pressure overload.

**Figure 3.**
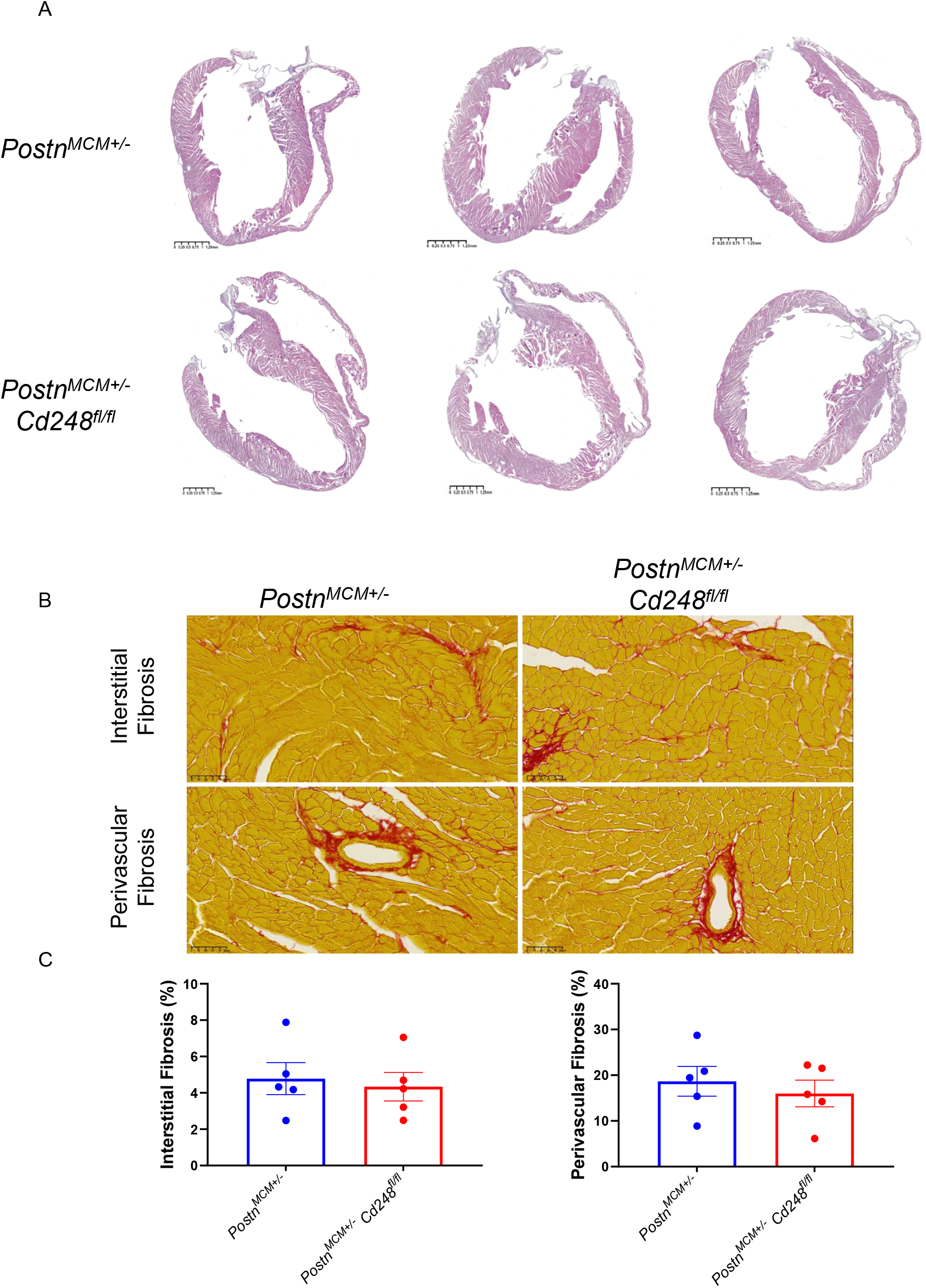
*Postn*^+^ myofibroblast-specific deletion of Cd248 does not alter cardiac fibrosis after pressure overload. (A) Representative low-magnification images of HE-stained long-axis heart sections 8 weeks after transverse aortic constriction (TAC). Scale bar, 1.25 mm. (B) High-magnification images of Picro-Sirius Red-stained heart section of control (Postn-MCM) and *Postn*^+^myofibroblast-specific Cd248 knockout (Cd248-cKO) mice are shown. Collagen is stained red, and myocardium appears yellow. Detailing interstitial fibrosis (top) and perivascular fibrosis (bottom) for each genotype were shown. Scale bar, 50 µm. (C, D) Quantification of the fibrotic area. Data are presented as mean ± SEM; n=5, each dot represents an individual animal. Statistical analysis was performed as described in the Methods.

**Figure 4.**
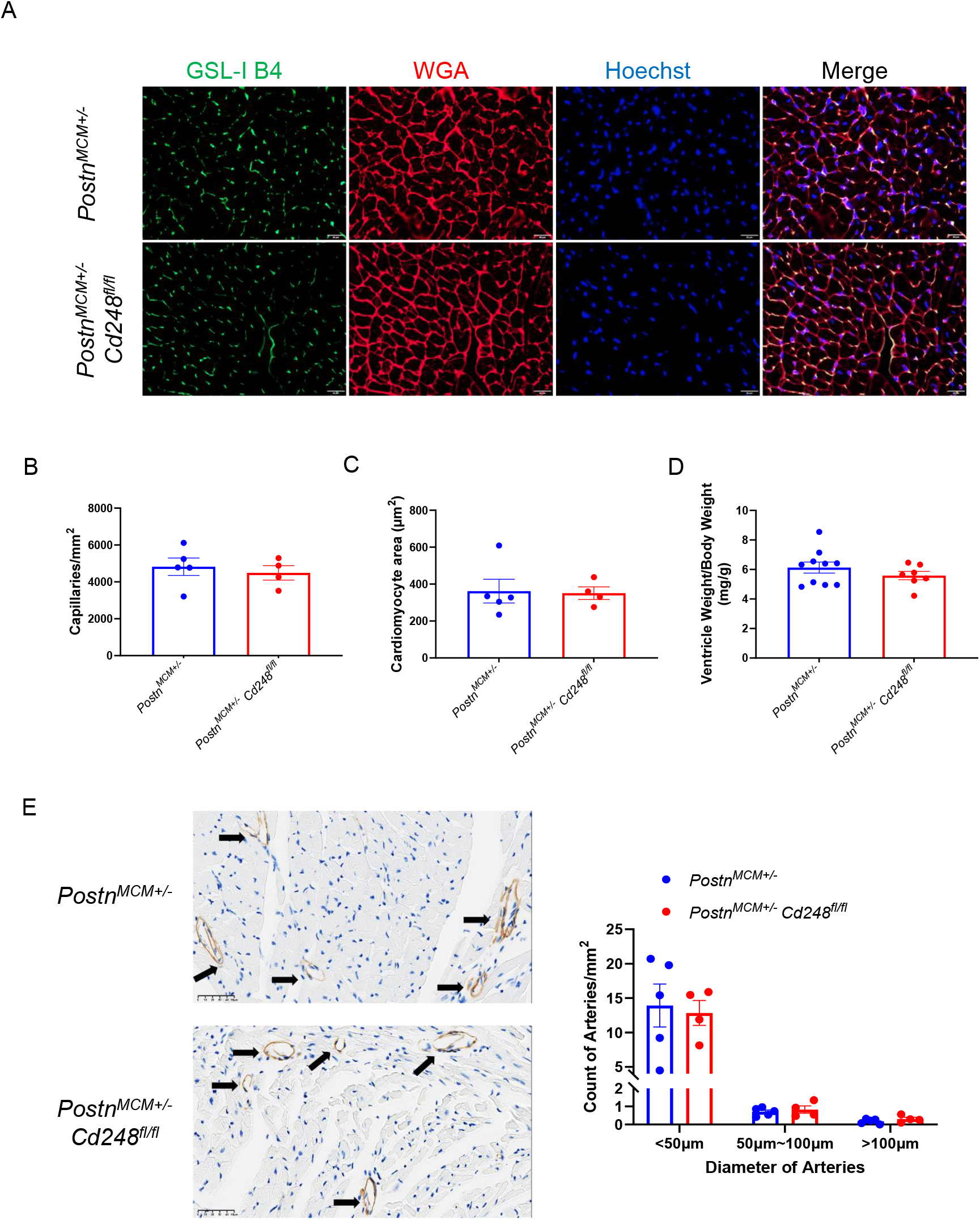
*Postn*^+^ myofibroblast-specific deletion of Cd248 does not alter cardiomyocyte hypertrophy or microvascular density in response to pressure overload. (A) Representative confocal images of left ventricular cross-sections from control and Cd248-cKO mice 8 weeks after TAC surgery. Capillaries are labeled with Isolectin-B4 (IB4, green), cardiomyocyte membranes with Wheat Germ Agglutinin (WGA, red), and nuclei with DAPI (blue). Scale bar, 50 µm. (B-D) Quantification of morphometric parameters. No significant differences were observed between control (blue bars) and Cd248-cKO (red bars) mice in (B) capillary density, (C) cardiomyocyte cross-sectional area, or (D) the ventricle weight to body weight ratio. (E) Representative images of arterioles identified by α-smooth muscle actin (α-SMA) immunohistochemistry (arrows). Scale bar, 50 µm. (F) Quantification of α-SMA^+^ arterioles, showing no significant differences in total arteriolar density (left) or in the distribution of arteriolar sizes (right) between genotypes. Data are presented as mean ± SEM; n=4-10, each dot represents an individual animal. Statistical analysis was performed as described in the Methods.

### Myofibroblast-Specific Deletion of *Cd248* Fails to Mitigate Cardiac Dysfunction

To confirm whether there would be changes in cardiac function or the cardiac muscle dynamic, which might be missed in histology, we performed echocardiographic assessment including traditional M-mode, B-mode pulsed-wave doppler, tissue doppler (Figure 5A) and speckle-tracking based muscle strain analysis in both groups. Consistent with histological findings, there are no changes in FS, EF, LVID, E/A and E/E’ at 8 weeks post-TAC between Postn-CD248 knockout and control mice (Figure 5B-G). Both groups developed significant and comparable concentric hypertrophy (Figure 5H and I). Speckle-tracking analysis based on strain analysis also revealed no difference between cardiac muscle movement between Postn-CD248 knockout and control mice (Supplemental Figure 5A-D).

**Figure 5.**
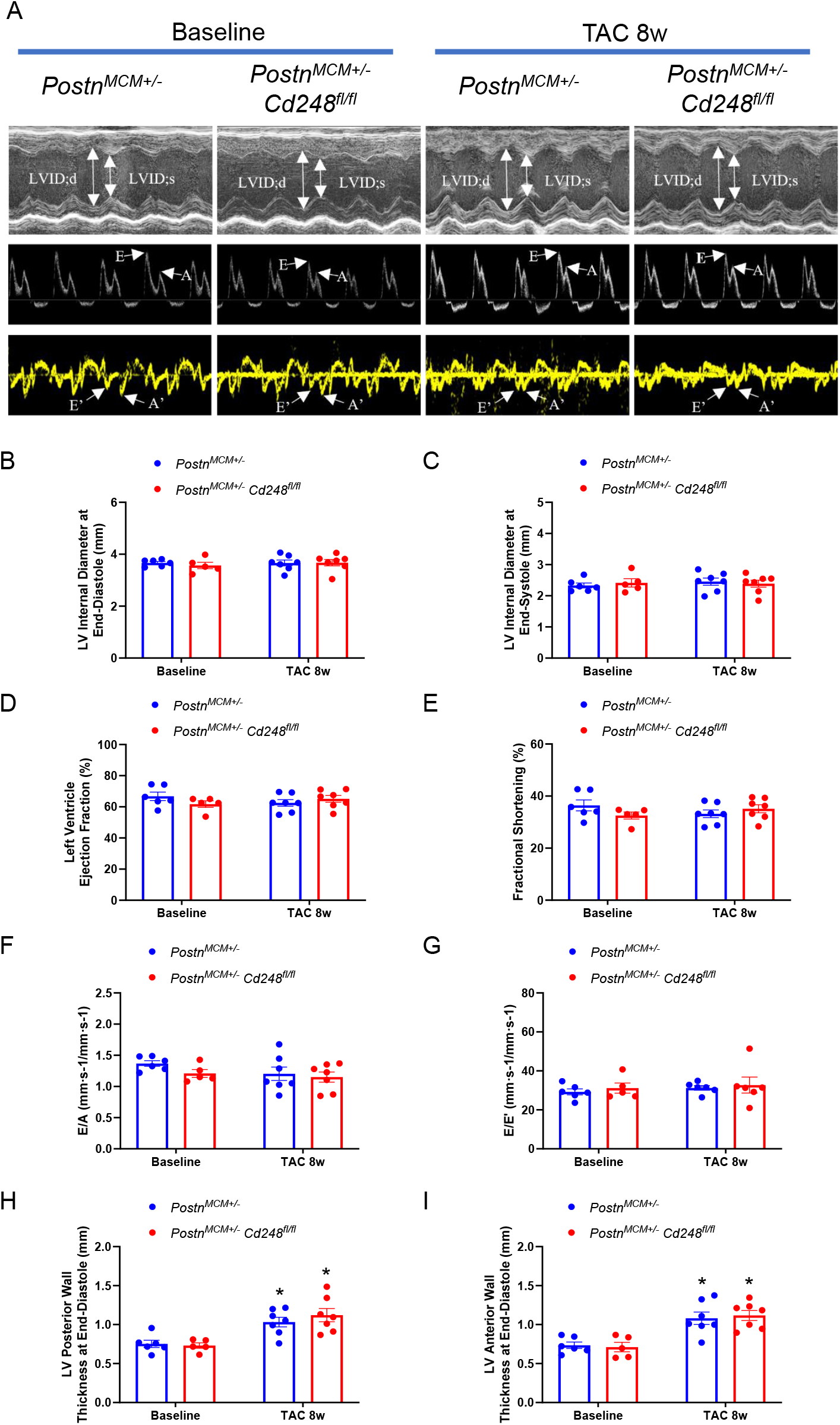
*Postn*^+^ myofibroblast-specific deletion of Cd248 does not alter cardiac function or dimensions after pressure overload. (A) Representative echocardiographic recordings from control and Cd248-cKO mice at baseline and 8 weeks after TAC. Images shown are M-mode, pulsed-wave Doppler of transmittal inflow, and tissue-Doppler imaging (TDI) of the mitral annulus. (B-I) Quantification of key echocardiographic parameters at baseline and 8 weeks post-TAC for control (blue bars) and Cd248-cKO (red bars) mice. Parameters shown include left ventricular internal dimensions at diastole and systole (LVID;d, LVID;s), systolic function (ejection fraction [EF], fractional shortening [FS]), diastolic indices (E/A, E’/A’, and E/E’) and posterior wall thickness (LVPW;d, LVPW;s). No significant differences were observed between genotypes. Data are presented as mean ± SEM; n=5-7, each dot represents an individual animal. Two-way Analysis of Variance (ANOVA) was performed, followed by Tukey’s multiple comparisons post-hoc test. \**P* < 0.05 vs. baseline.

### Single-Cell Analysis Reveals Distinct Transcriptional Programs in *Cd248*^+^ and *Postn*^+^ Fibroblast Subpopulations

To understand the potential heterogeneity of the target cell population, we further analyzed the fibroblast cluster from the scRNA-seq data of TAC hearts^[15]^. Unsupervised clustering resolved the fibroblasts into 11 distinct subpopulations (Figure 6A). A bivariate feature plot in UMAP revealed four main groups based on the expression of Cd248 and Postn: double-positive (*Cd248*^+^*Postn*^+^), *Cd248* single-positive, *Postn* single-positive, and double-negative cells (Figure 6B). To elucidate the functional differences between these subpopulations, we performed differential gene expression analysis followed by Gene Ontology (GO) enrichment. When comparing the double-positive (*Cd248*^+^*Postn*^+^) population to the *Postn* single-positive (*Postn*^+^*Cd248*^−^) population (Figure 6C), we found that the double-positive cells were significantly enriched for genes associated with chemotaxis and migration. Importantly, when comparing the double-positive population to the Cd248 single-positive (*Postn*^−^*Cd248*^+^) population, the enriched terms were related to extracellular matrix/structure organization (Figure 6D). These results suggest that fibroblasts co-expressing *Cd248* and *Postn* represent an immune reaction related subpopulation, while the fibrosis associated effects might be dependent on the *Postn*^−^*Cd248*^+^ fibroblast subsets which is not targeted in *PostnMCM*+*/-;Cd248fl/fl* mouse model in the pressure-overloaded heart.

**Figure 6.**
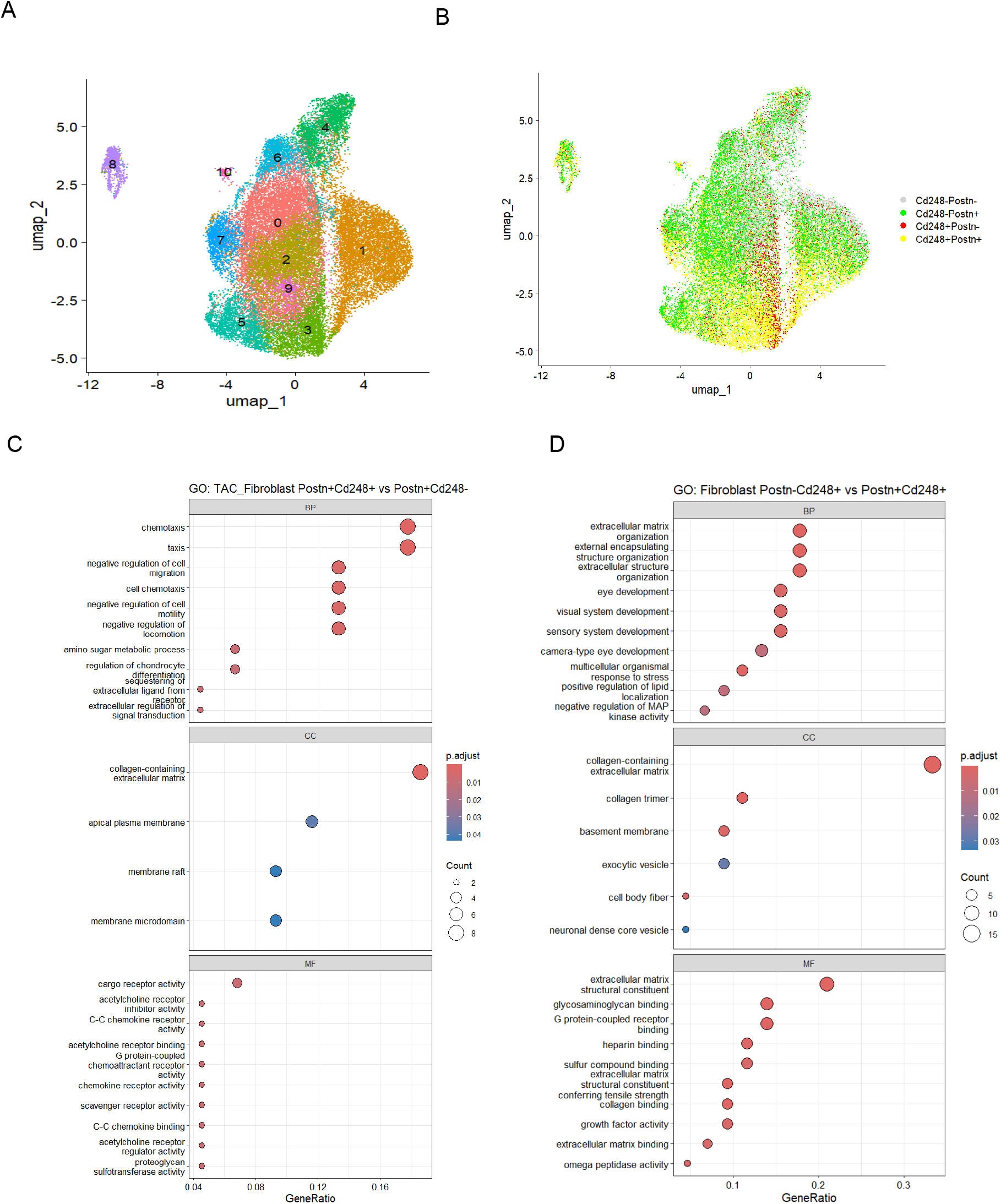
Single-cell transcriptomic analysis reveals distinct biological programs in fibroblast subpopulations from pressure-overloaded hearts. (A) UMAP visualization of cardiac fibroblasts isolated from mice 14 days after TAC, showing unsupervised clustering into distinct subpopulations (0–10). (B) *Cd248*^*-*^*Postn*^*-*^ (grey), *Cd248*^*-*^*Postn*^+^ (green), *Cd248*^+^*Postn*^*-*^ (red)and their co-expression (yellow) within the fibroblast UMAP. (C-D) Gene Ontology (GO) enrichment analysis of differentially expressed genes (DEGs) from the TAC fibroblast population. Dot plots show the top enriched biological programs comparing (C) *Postn*^+^*Cd248*^+^ versus *Postn*^+^*Cd248*^−^ fibroblasts and (D) *Postn*^*-*^*Cd248*^+^ versus *Postn*^+^*Cd248*^+^ fibroblasts. Dot size corresponds to the number of DEGs mapped to each term, and color indicates the false discovery rate (FDR). (BP, Biological Process; CC, Cellular Component; MF, Molecular Function).

## Discussion

The primary finding of this investigation is the absence of an observable cardiac phenotype following the specific deletion of Cd248 in *Postn*-expressing myofibroblasts within a murine model of pressure-overload cardiomyopathy. This result is scientifically significant as it indicates that the pro-fibrotic function of Cd248, which has been well-documented in the context of ischemic cardiac injury, may not be conserved across different etiologies of heart disease. Specifically, it suggests a context-dependent role for Cd248 within the *Postn*^+^ myofibroblast lineage.

The rationale for targeting the *Postn*^+^ cell lineage was based on extensive prior validation. This approach has been repeatedly validated as an effective strategy to interrogate fibroblast function in the pressure-overloaded heart in our hands as well as in other labs^[2-5, 7, 8]^. For example, Takeda et al. used a *Postn-Cre* driver to show that fibroblast-specific deletion of KLF5 attenuates both fibrosis and cardiomyocyte hypertrophy by disrupting paracrine IGF-1 signaling after TAC^[6]^. Similarly, our group and others have used *Postn-MerCreMer* mice to demonstrate that specifically inhibiting the canonical TGFβ or WNT/β-catenin signaling as well as collagen chaperone Hsp47 in activated fibroblasts robustly blocked pressure-overload-induced fibrosis and secondarily attenuated the hypertrophic response^[3, 7, 8]^. Collectively, these studies confirmed that targeting key molecular pathways within the *Postn*+ fibroblast population is a valid and effective strategy for mitigating pressure-overload pathology.

The hypothesis for the present study was further supported by recent findings that identified Cd248 as a specific marker for a late-activated, pathological fibroblast subpopulation in ischemic injury models. Both Chen et al. (2025) and Li et al. (2025) independently showed that therapeutic interventions targeting these *Cd248*^+^ cells successfully reduced fibrosis and improved cardiac function after myocardial infarction or ischemia/reperfusion^[13, 14]^. Our re-analysis of published single-cell sequencing data confirms that *Cd248*^+^ fibroblasts constitute a substantial portion (over 30%) of the total fibroblast population in the pressure-overloaded heart, suggesting this subpopulation should have a meaningful impact on pathology. The logical synthesis of these prior findings—the established role of the *Postn*^+^ cell target in TAC and the validated function of Cd248 in ischemia—created a strong expectation of efficacy for our experiment.

The null phenotype reported here stands in notable contrast to these positive findings and is therefore highly informative. This discrepancy suggests that the function of Cd248 on *Postn*^+^ fibroblasts is profoundly context-dependent, differing between ischemic injury and chronic pressure overload. While Cd248 appears crucial for the pathological activity of fibroblasts in post-infarction remodeling, our data demonstrate it is dispensable within the same cell lineage when the injury stimulus is sustained hemodynamic stress. This highlights the distinct biological mechanisms underlying different fibrotic diseases and cautions against the assumption that anti-fibrotic targets are universally applicable.

The absence of a phenotype following Cd248 deletion in *Postn*^+^ cells suggests that the critical functions of Cd248 in the pressure-overload model may be mediated by other, Postn-negative cell populations. Our single-cell analysis confirms that Cd248 is also highly and persistently expressed in *Pdgfrβ*^+^ mural cells, indicating that pericytes are also a major, constitutive source of Cd248 in the TAC heart. These cells may therefore contribute to the development of pathology in pressure-overload hearts. Future studies employing pericyte-specific lineage-tracing and gene deletion are required to fully elucidate the role of pericyte-derived CD248 in pressure-overload induced cardiac injury.

From a translational perspective, targeting CD248 with neutralizing antibodies has shown considerable promise as an anti-fibrotic strategy in a range of preclinical settings. Efficacy has been demonstrated not only in animal models of ischemic heart injury^[13, 14]^ but also in fibrotic conditions of the kidney^[18]^, liver^[19]^, and skin^[20]^. This body of evidence, combined with the favorable safety profile of the CD248-targeting antibody Ontuxizumab in oncology trials^[10]^, has generated significant interest in its potential application for non-malignant fibrotic diseases. However, our findings introduce a critical note of caution. The lack of efficacy in the pressure-overload model, despite the positive results in other pathologies, underscores that our understanding of CD248’s function is incomplete. Therefore, a better and more in-depth investigation of the context-dependent role of CD248 in various pathological conditions is critical to successfully guide and refine the clinical translation of these promising anti-fibrotic therapies.

## Conclusion

In summary, our findings indicate that the specific ablation of CD248 in the Postn+ myofibroblast lineage does not substantially mitigate fibrosis or cardiac dysfunction in this murine model of pressure-overload cardiomyopathy. These results highlight the context-dependent function of CD248, suggesting its role is not universally conserved across different etiologies of cardiac injury. Therefore, the future development of anti-fibrotic therapies targeting this molecule should consider the specific disease model and may need to address its function in other cell types, such as pericytes, to achieve therapeutic efficacy.

## Data availability

Datasets generated during and/or analysed during the current study are available from J.X. on reasonable request. Source data are provided with this paper.

## Acknowledgements and funding

We thank the staff of the scientific core facility the First Affiliated Hospital of Sun Yat-sen University for technical assistance. This study was supported by Natural Science Foundation of China (82270274 to F.-l.X., 32371199 to J.X.,), Guangdong Provincial Science and Technology Department (2023ZT10Y154 to F.-l.X.), Chinese National Key Research and Development Program (no. 2023YFC2307004 to F.-l.X.).

## Author contributions

J.X., F.-l.X. and C.L. conceived the project and wrote the manuscript with the help from all of the other authors. D.L., D.Z., T.T. and Z.H. designed and performed the experiments, with assistance from the other authors. M.W., Y.L. and Y.Z. assisted in carrying out experiments. F.-l.X. and J.X. supervised the project. D.L., D.Z., T.T. and Z.H. contributed equally to this study. J.X. is the lead contact.

---

**Supplemental Figure 1.** Single-cell analysis reveals enriched CD248 expression in pericytes and fibroblasts from human hypertrophic cardiomyopathy hearts.

(A, B) UMAP visualization of cardiac cells from healthy control and hypertrophic cardiomyopathy (HCM) human hearts, color-coded by cell type (A) or sample group (B). (C) UMAP feature plot showing normalized CD248 expression across all cells. (D) Bar plot quantifying the percentage of CD248^+^ cells within each cell lineage.

**Supplemental Figure 2.** Cd248 mRNA expression is upregulated in murine models of cardiac injury.

(A) Relative mRNA expression of Cd248 across various organs from healthy adult C57BL/6J mice, demonstrating the highest basal expression in the heart, lung, and kidney. (B) Myocardial Cd248 mRNA expression is significantly increased 1 week after pressure overload induced by transverse aortic constriction (TAC) compared to sham-operated controls. (C) Myocardial Cd248 mRNA expression is significantly increased in the infarct border zone 3 days after myocardial infarction (MI) compared to sham-operated controls. Data are presented as mean ± SEM; n=3–6 mice per group. \**P* < 0.05 vs. sham, determined by an unpaired Student’s *t-test*.

**Supplemental Figure 3.** Cd248 knockdown alters the expression of genes related to myofibroblast activation and extracellular matrix remodeling.

Relative mRNA expression of *Cd248* (A), *Col3a1* (B), *Col1a1* (C), and *Postn* (D) in cultured cardiac fibroblasts transfected with control siRNA (si-Scrambled) or siRNA targeting Cd248 (si-Cd248) for 48 hours, followed by stimulation with TGF-β1 (5 ng/mL) for 24 hours. Data are presented as mean ± SEM; n=6 independent experiments per group. Two-way Analysis of Variance (ANOVA) was performed, followed by Tukey’s multiple comparisons post-hoc test. \**P* < 0.05 vs. si-Scrambled control; #*P* < 0.05 vs. si-Scrambled with TGF-β1 stimulation.

**Supplemental Figure 4.** Validation of the TAC model and tamoxifen-inducible, periostin-lineage tracing of activated fibroblasts.

(A) Representative pulse-wave Doppler tracings confirming successful transverse aortic constriction (TAC). After surgery, peak velocity across the aortic arch increased from ∼1 m/s to over 3.5 m/s, with a corresponding increase in the trans-stenotic pressure gradient, indicating a severe and consistent stenosis. (B) Picro-Sirius red staining of left ventricular sections demonstrates robust induction of interstitial and perivascular fibrosis (collagen stained red) in hearts subjected to TAC for 6 weeks compared to sham-operated controls. Scale bars as indicated. (D) The tamoxifen (TAM)-inducible *Postn-MerCreMer* (*Postn-MCM*) lineage tracing strategy, employing a Rosa26-LSL-tdTomato reporter allele was used. Representative fluorescence images from *Postn-MCM*;LSL-tdTomato hearts showing the progressive accumulation of tdTomato-positive activated fibroblasts (red) in the interstitium and perivascular space at 2 and 6 weeks post-TAC. Nuclei are counterstained with Hoechst (blue). Scale bars as indicated.

**Supplemental Figure 5.** Post-operative survival and aortic peak velocity in *Postn*^+^ myofibroblast-specific Cd248 deletion mice.

(A) Kaplan-Meier survival analysis over an 8-week period following TAC surgery. There was no significant difference in survival between the control (blue line; n=11) and Cd248-cKO mice (red line; n=8), as determined by the log-rank test. (B) Quantification of aortic peak velocity by Doppler echocardiography 2 weeks after transverse aortic constriction (TAC) surgery. The data show a comparable degree of stenosis and pressure overload in both Cd248-cKO mice and their littermate controls. Each dot represents an individual animal; bars indicate mean ± SEM, n=7-11.

**Supplemental Figure 6.** *Postn*^+^ myofibroblast-specific deletion of Cd248 does not alter myocardial deformation in response to pressure overload.

Speckle-tracking echocardiography was performed at baseline and 8 weeks after transverse aortic constriction (TAC). Control littermates (blue bars) are compared with Cd248-cKO (red bars). (A, B) Quantification of global radial strain rate (A) and reverse global radial strain rate (B). (C, D) Quantification of global longitudinal strain (C) and reverse global longitudinal strain (D). No significant differences in any strain parameter were observed between genotypes at either baseline or 8 weeks post-TAC. Data are presented as mean ± SEM; n=5-7, each dot represents an individual animal. Two-way Analysis of Variance (ANOVA) was performed, followed by Tukey’s multiple comparisons post-hoc test. \**P* < 0.05 vs. baseline.

